# Large mRNA language foundation modeling with NUWA for unified sequence perception and generation

**DOI:** 10.1101/2025.11.01.686058

**Authors:** Yunshan Zhong, Weiqi Yan, Yiming Zhang, Kuanhuan Tan, Yutaka Saito, Bian Bian

## Abstract

The mRNA serves as a crucial bridge between DNA and proteins. Compared to DNA, mRNA sequences are much more concise and information-dense, which makes mRNA an ideal language through which to explore various biological principles. In this study, we present NUWA, a large mRNA language foundation model leveraging a BERT-like architecture, trained with curriculum masked language modeling and supervised contrastive loss for unified mRNA sequence perception and generation. For pretraining, we utilized large-scale mRNA coding sequences comprising approximately 80 million sequences from 19,676 bacterial species, 33 million from 4,688 eukaryotic species, and 2.1 million from 702 archaeal species, and pre-trained three domain-specific models respectively. This enables NUWA to learn coding sequence patterns across the entire tree of life. The fine-tuned NUWA demonstrates strong performance across a variety of downstream tasks, excelling not only in RNA-related perception tasks but also exhibiting robust capability in cross-modal protein-related tasks. On the generation front, NUWA pioneers an entropy-guided strategy that enables BERT-like models in generating mRNA sequences, producing natural-like sequences that accurately recapitulate species-specific codon usage patterns. Moreover, NUWA can be effectively fine-tuned on small, task-specific datasets to generate functional mRNAs with desired properties, including sequences that do not exist in nature, and to design coding sequences for diverse proteins in biomanufacturing, vaccine development, and therapeutic applications. To our knowledge, NUWA represents the first mRNA language model for unified sequence perception and generation, providing a versatile and programmable platform for mRNA design.

## 1 Introduction

mRNA plays a crucial role in the central dogma by conveying genetic information from DNA to protein [1, 2]. Unlike genomic DNA, which harbors introns, enhancers, and a variety of non-coding regulatory elements, mature mRNA is assembled from spliced exons and comprises a 5’ UTR, a protein-coding sequence (CDS) region, and a 3’ UTR [3]. This more “focused” sequence grammar, together with layered regulatory signals (e.g., codon bias and UTR structure), enhances the signal-to-noise ratio, making mRNA an ideal substrate for sequence modeling and for using language models to uncover biologically plausible patterns [4, 5].

The remarkable success of mRNA-based vaccines during recent global health crises has dramatically underscored the tremendous therapeutic potential of mRNA technology, heralding a new era in vaccinology and precision medicine [1, 6, 7]. However, despite this transformative promise, the efficient and rational design of highly functional mRNA sequences remains a formidable challenge at the intersection of computational biology and biopharmaceutical engineering [8].

A central challenge lies in the multifaceted nature of mRNA sequence optimization, which requires the simultaneous consideration of multiple factors, including GC content modulation for RNA stability and immunogenicity control, and the avoidance of cryptic regulatory elements such as splicing signals or internal ribosome entry sites (IRES) that may compromise predictable expression. [9–11]. These challenges are further compounded by codon degeneracy, an evolutionarily advantageous feature for biodiversity that nonetheless often impedes efficient heterologous expression [12–14]. Without meticulous host-specific optimization, foreign gene sequences may suffer from suboptimal translation kinetics, resulting in ribosomal stalling, premature termination, or amino acid misincorporation, ultimately leading to reduced protein yields, improper folding, loss of functional activity, or even cytotoxic effects [12, 15].

Traditionally, addressing these complex and often competing design requirements has relied heavily on experimental trial-and-error approaches [9, 16]. Conventional computational strategies, such as codon adaptation index (CAI) maximization or frequency matching, remain largely heuristic and labor-intensive [14, 17]. These methods typically focus on localized sequence features, such as individual codon replacements or short-range GC content, while overlooking higher-order regulatory elements and global structural context of mRNA sequence, which are increasingly recognized as critical determinants of expression efficiency [12, 15]. This limitation significantly constrains the rapid development of mRNA therapeutics.

Large language models have revolutionized natural language processing by capturing long-range dependencies and semantic representations [18, 19]. Their success has inspired analogous efforts in biology: protein language models (e.g., the ESM series) and DNA language models (e.g., the Evo series) have harnessed enormous sequence corpora to excellent effect [20–22]. While RNA-focused models such as RNA-FM for non-coding RNAs and CodonBERT/CodonTransformer for targeted mRNA tasks and a few organisms, and also PlantRNA-FM, which specifically focuses on plant species have emerged [5, 23–25].mRNA language models, trained on extensive multi-species mRNA sequences, offer a transformative approach to this longstanding mRNA design problem. By learning from mRNA sequence landscapes across diverse organisms, these advanced models capture not only host-specific codon usage preferences but also complex, context-dependent patterns such as RNA secondary structure, splicing signals, internal ribosome entry sites, and immunogenic motifs [23, 26]. This comprehensive understanding enables the generation of fully optimized mRNA sequence that strategically balance translational efficiency with structural stability and minimal immunogenicity [27]. Consequently, mRNA language models provide a powerful and scalable computational framework for the *de novo* design of synthetic mRNA sequences tailored for robust expression in desired host systems, thereby significantly accelerating the development of next-generation mRNA-based therapeutics and recombinant protein production platforms [24, 27]. However, comprehensive large mRNA language foundation models that unify both sequence perception and generative capabilities, trained on large-scale datasets across multiple species, remain scarce.

In this study, we present NUWA, the largest mRNA language model that was pretrained on mRNA coding sequences. NUWA leverages a BERT-like architecture, trained with curriculum masked language modeling and supervised contrastive loss, to achieve strong performance across a variety of mRNA sequence perception and generation tasks. To capture codon usage patterns across the tree of life, we pre-trained three domain-specific mRNA language models on large-scale datasets comprising approximately 80 million sequences from 19,676 bacterial species, 33 million from 4,688 eukaryotic species, and 2.1 million from 702 archaeal species. Furthermore, for the mRNA perception task, we fine-tuned this model on a curated dataset for many RNA property-related profiles, including Structure-function-engineering suite tasks. In addition, our model exhibits generative ability, which offers valuable insights into the data-driven design of stable and efficient mRNA sequence.

## 2 Results

### 2.1 NUWA unified mRNA sequence perception and generation

NUWA incorporates mRNA coding sequence from the three domains of life (Figure 1.A)and unifies mRNA sequence perception and generation through an encoder-only architecture (Figure 1.B) that learns bidirectional, context-complete representations while being explicitly regularized for both discriminative perception and generative fidelity. Unlike encoder-decoder or decoder-only models that rely on unidirectional auto-regression and thus inherit directional biases, NUWA’s BERT-style encoder ingests the entire codon sequence at once, enabling it to capture subtle, non-local constraints that often govern regulatory logic in mRNA, for example, how a downstream motif can shape the optimal choice of an upstream codon. This bidirectional, holistic view is essential for decoding the intertwined structural, functional, and regulatory dependencies that span local motifs and long-range interactions.

**Fig. 1:**
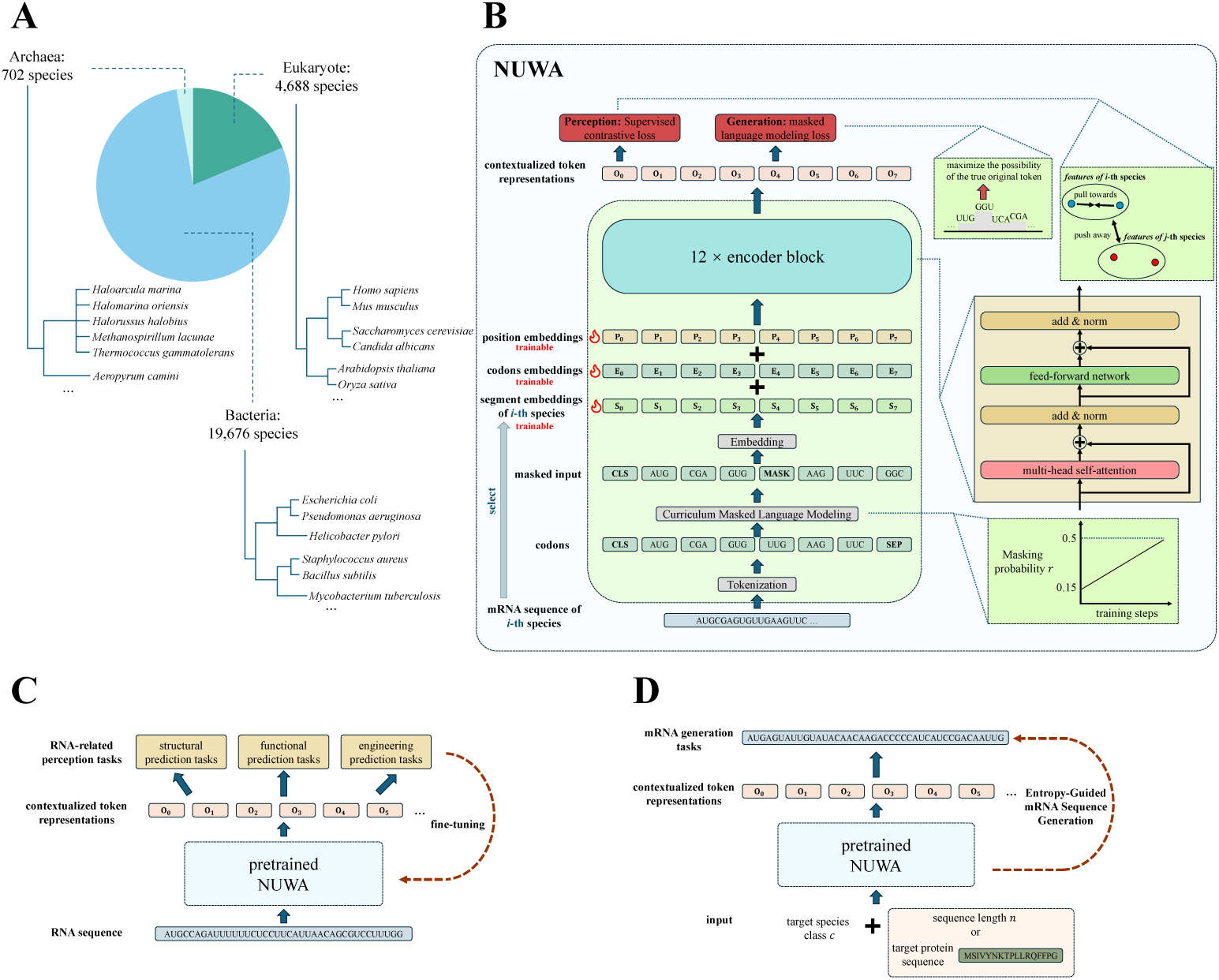
The overview of NUWA. (A) Taxonomic distribution of the pretraining dataset, comprising mRNA coding sequences from three domains of life, including bacteria, archaea, and eukaryotes, covering 19,676, 702, and 4,688 species, respectively. (B) Schematic of the NUWA architecture and pretraining strategy, which integrates supervised contrastive learning (SCL) for perception tasks and curriculum masked language modeling (C-MLM) for generative objectives. (C) Fine-tuning of the pretrained NUWA model on diverse RNA-related perception tasks, including structural, functional, and engineering predictions. (D) Entropy-guided mRNA sequence generation pipeline using pretrained NUWA to produce biologically plausible, organism-specific mRNA sequences.

At the input level, NUWA adopts codon-granular tokenization (64 codons tokens + special tokens), preserving the genetic code’s natural compositional unit and improving the signal-to-noise ratio for both perception and generation. Species-specific segment embeddings further condition the encoder on archaea, bacteria, or eukaryote domains (Figure 1.A), acknowledging the deep evolutionary divergence in codon usage, structure, and regulation. Practically, NUWA is instantiated as three specialized encoders, one for each domain of life, so that each learns features tuned to its biological constraints while sharing the same modeling principles and objectives.

Two complementary pretraining objectives fuse perception and generation within the encoder-only model. First, Supervised Contrastive Loss (SCL) shapes a discriminative representation space: samples from the same class are pulled together while those from different classes are pushed apart. This enforces clusterable, semantically meaningful embeddings (via [CLS]) that benefit downstream classification, ranking, or retrieval-style tasks (Figure 1.C), i.e., “perception”. Second, Curriculum Masked Language Modeling (C-MLM) progressively increases the masking ratio from an easier regime (e.g., 15%) to a challenging one (e.g., 50%). Early in training, abundant context encourages stable semantic grounding; later, sparse context compels the model to infer larger spans, strengthening its ability to reconstruct plausible sequences under severe information occlusion (Figure 1.D), i.e., “generation”. The curriculum, therefore, transitions NUWA from learning fine-grained local semantics to mastering global sequence infill, ensuring the encoder’s representations remain useful for both perception and generation.

As shown in Figure 1.B, architecturally, multi-head self-attention layers (with residuals, layer norm, and position embeddings) compute rich, bidirectional token states that serve as a universal substrate. Because generation is trained as masked reconstruction rather than left-to-right prediction, NUWA can honor bidirectional constraints at inference time for tasks such as codon infilling, constrained editing, and motif-aware redesign, without abandoning the same encoder used for perception tasks. This encoder-only design yields a compact, data-efficient solution: one backbone, two objectives, and unified representations that transfer across structural prediction, functional inference, and engineering-oriented design.

To evaluate the representational power of our pretrained model, we projected the embedding vectors of different archaeal, bacterial, and eukaryotic sequences using UMAP (Figure S1.A-S1.C). Sequences from the same species formed compact clusters, while those from different species were well separated, indicating that NUWA learned meaningful inter-species distinctions in the embedding space. Leveraging this organism-aware representation, NUWA can generate mRNA coding sequences specific to a given species. We can even generate mRNA sequences that are optimized for expression in that species but do not exist in nature, enabling the construction of a synthetic mRNA database tailored to any target organism.

In summary, NUWA is a codon and organism-aware, domain-specialized, encoderonly framework, augmented with SCL for discriminative structure learning and CMLM for reconstructive pressure, which provides a principled route to unify sequence perception and generation within a single model.

### 2.2 Fine-tuned NUWA reveals superior performance in diverse RNA-related tasks

To assess NUWA on downstream RNA tasks (spanning RNA function, RNA engineering, and RNA structure), we assembled benchmarks and evaluation protocols from the CodonBERT and BEACON papers and reproduced the settings for fair comparison (Tables 1 and 3).

**Table 1:** Comparison of NUWA to prior methods on six downstream tasks.

| Model | mRFP<br>expression | Fungal<br>expression | *E. coli*<br>proteins | mRNA<br>stability | Tc-riboswitch | SARS-CoV-2<br>vaccine<br>degradation |
| --- | --- | --- | --- | --- | --- | --- |
| Nucleotide-based |  |  |  |  |  |  |
| Plain TextCNN | 0.62 | 0.53 | 0.39 | 0.01 | 0.41 | 0.55 |
| RNABERT+TextCNN | 0.40 | 0.41 | 0.39 | 0.16 | 0.47 | 0.64 |
| RNA-FM+TextCNN | 0.80 | 0.59 | 0.43 | 0.34 | 0.58 | 0.74 |
| Codon-based |  |  |  |  |  |  |
| TF-IDF | 0.57 | 0.68 | 0.44 | **0.54** | 0.49 | 0.69 |
| Plain TextCNN | 0.78 | 0.76 | 0.36 | 0.26 | 0.43 | 0.80 |
| Codon2vec+TextCNN | 0.77 | 0.61 | 0.43 | 0.33 | 0.56 | 0.70 |
| CodonBERT | 0.85 | **0.88** | 0.55 | 0.51 | 0.56 | 0.77 |
| NUWA | **0.89** | 0.81 | **0.61** | 0.53 | **0.62** | **0.83** |

**Table 2:** Overview of task datasets used in Table 1. “Cls” indicates “classification”, “Spr” indicates spearman correlation coefficient, and “ACC” indicates accuracy.

| Dataset | Metric | Task<br>Type | #Train<br>#Validation<br>#Test | Max/Mean<br>Length |
| --- | --- | --- | --- | --- |
| mRFP Expression | Spr | Regression | 1,021/219/219 | 678/678 |
| Fungal expression | Spr | Regression | 5,089/1,000/1,000 | 3,000/1195 |
| *E. coli* proteins | ACC | Multi-class Cls | 4,348/1,000/1,000 | 3,000/817.8 |
| Tc-Riboswitches | Spr | Regression | 248/53/54 | 75/71.9 |
| mRNA stability | Spr | Regression | 45,749/9,803/9,804 | 3,000/1,342.2 |
| SARS-CoV-2 Vaccine Degradation | Spr | Regression | 1,600/400/400 | 81/81 |

**Table 3:** Comparison of NUWA to prior methods on ten downstream tasks.

| Model | SSP | CMP | DMP | APA | NcRNA | Modif | MRL | PRS | CRI-On | CRI-Off |
| --- | --- | --- | --- | --- | --- | --- | --- | --- | --- | --- |
| Nucleotide-based |  |  |  |  |  |  |  |  |  |  |
| RNA-FM | **0.69** | 0.48 | 0.51 | 0.70 | **0.97** | **0.95** | 0.79 | 0.56 | 0.32 | 0.02 |
| RNABERT | 0.57 | 0.45 | 0.48 | 0.58 | 0.69 | 0.83 | 0.30 | 0.55 | 0.30 | 0.04 |
| RNA-MSM | 0.58 | 0.57 | 0.37 | 0.70 | 0.85 | **0.95** | 0.83 | 0.57 | 0.35 | 0.04 |
| Splice-H510 | 0.65 | 0.46 | 0.56 | 0.59 | 0.96 | 0.63 | 0.83 | 0.55 | 0.27 | 0.04 |
| Splice-MS510 | 0.43 | 0.53 | 0.10 | 0.52 | 0.96 | 0.56 | **0.85** | 0.51 | 0.27 | 0.03 |
| Splice-MS1024 | 0.68 | 0.47 | 0.56 | 0.60 | 0.96 | 0.53 | 0.67 | 0.58 | 0.28 | 0.05 |
| UTR-LM-MRL | 0.60 | 0.46 | 0.55 | 0.65 | 0.90 | 0.56 | 0.78 | 0.57 | 0.28 | 0.04 |
| UTR-LM-TE&EL | 0.60 | 0.60 | 0.55 | **0.72** | 0.81 | 0.60 | 0.83 | 0.53 | 0.32 | 0.03 |
| BEACON-B | 0.64 | 0.60 | 0.56 | 0.71 | 0.95 | **0.95** | 0.72 | 0.55 | 0.26 | 0.04 |
| BEACON-B512 | 0.59 | 0.61 | 0.57 | **0.72** | 0.95 | **0.95** | 0.72 | 0.55 | 0.28 | 0.04 |
| Codon-based |  |  |  |  |  |  |  |  |  |  |
| UTRBERT | 0.60 | 0.51 | 0.51 | 0.70 | 0.93 | **0.95** | 0.84 | 0.57 | 0.30 | 0.04 |
| NUWA | 0.65 | **0.63** | **0.58** | **0.72** | 0.95 | **0.95** | 0.84 | **0.59** | **0.38** | **0.07** |

Across six representative tasks in expression and stability (Table 1), NUWA achieves the best scores on mRFP expression (0.89), *E. coli* proteins (0.61), Tc-riboswitch (0.62), and SARS-CoV-2 vaccine degradation (0.83), outperforming CodonBERT and other baselines. CodonBERT is slightly better on Fungal expression (0.88 vs. 0.81), and TF-IDF edges NUWA on mRNA stability (0.54 vs. 0.53) under this setup. Overall, codon-based approaches outperform nucleotide-level methods, and NUWA is consistently the strongest codon-level model, indicating more faithful modeling of codon usage bias, context dependencies, and regulatory constraints.

For the structure–function–engineering suite in BEACON’s ten tasks(Table 3), NUWA (codon-based) surpasses UTRBERT on 8/10 tasks and ties or remains comparable on the rest.

Structure: NUWA attains 0.65 (SSP), 0.63 (CMP), and 0.58 (DMP), exceeding UTRBERT (0.60/0.51/0.51), highlighting better capture of local-to-long-range constraints relevant to secondary/tertiary organization.

Function: NUWA reaches 0.72 (APA), 0.95 (NcRNA), 0.95 (Modif), and 0.84 (MRL),matching or surpassing UTRBERT, demonstrates robust sensitivity to poly(A) usage, ncRNA functional signals, RNA modifications, and translation efficiency.

Engineering: NUWA outperforms on PRS (0.59), CRI-On (0.38), and CRI-Off (0.07) versus UTRBERT (0.57/0.30/0.04), showing stronger transfer to programmable RNA switches and CRISPR on/off-target prediction.

Taken together, these results show NUWA achieves state-of-the-art or competitive performance on 4/6 expression/stability tasks and 8/10 structure–function–engineering tasks in the codon-based setting. Coupled with our representation analyses, this indicates that NUWA not only perceives and captures mRNA sequence patterns across evolutionary lineages but also translates those representations into broad downstream gains, consistent with a unified, encoder-only design that supports the strong perception ability.

### 2.3 NUWA generates natural-like mRNA sequence

To evaluate the generative capability of NUWA, we developed an entropy-guided mRNA sequence generation pipeline (Figure 2.A). During generation, positions with the highest token-level entropy were iteratively selected and filled by sampling from the model’s predicted codon distribution, enabling efficient exploration of sequence space while maintaining biological plausibility. This strategy ensures that NUWA produces diverse yet functionally consistent mRNA sequences conditioned on the target protein and host species.

**Fig. 2:**
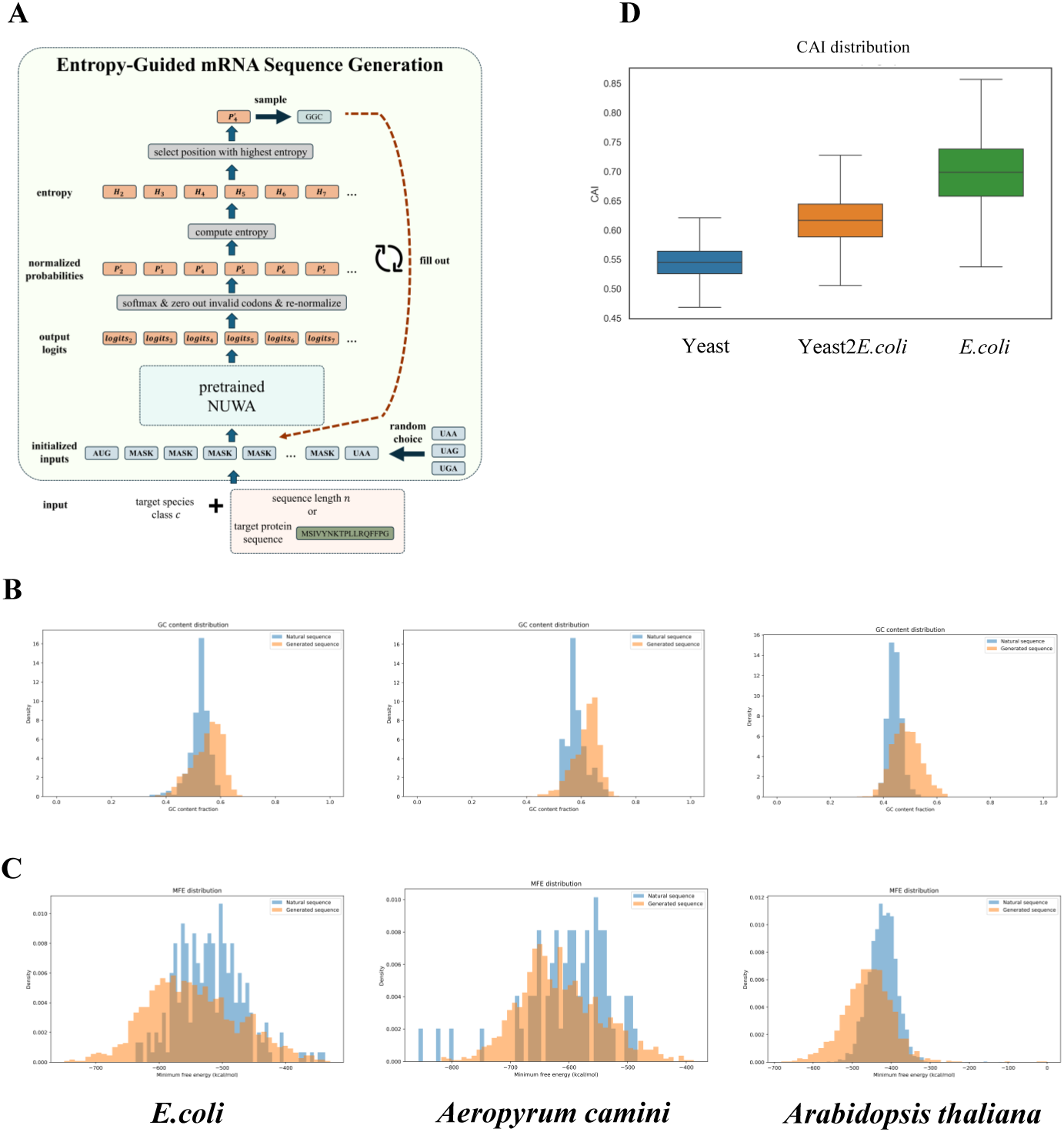
Entropy-guided mRNA sequence generation and comparative analysis. (A) Overview of the entropy-guided mRNA sequence generation pipeline based on the pretrained NUWA model. (B) GC-content distribution of generated mRNA coding sequences compared with natural mRNA coding sequences across representative species. (C) Minimum free-energy (MFE) distribution of generated versus natural mRNA coding sequences across representative species. (D) Codon adaptation index (CAI) comparison among yeast native mRNA, yeast-optimized mRNA sequences expressed in *E. coli* generated by NUWA, and endogenous *E. coli* mRNA sequences.

We next compared several intrinsic features of the generated sequences with those of natural mRNAs. As shown in Figure 2.B, the GC-content distributions of the generated mRNAs closely matched those of natural sequences across representative organisms, including *E. coli, Archaeoglobus camini*, and *Arabidopsis thaliana*, indicating that NUWA preserves native-like compositional biases. Similarly, the minimum free energy (MFE) distributions of generated sequences were nearly indistinguishable from those of their natural counterparts (Figure 2.C), suggesting that the model captures relevant structural determinants underlying RNA stability.

Finally, we assessed codon-level optimization using the codon adaptation index (CAI) (Figure 2.D). Generated sequences optimized for *E. coli* exhibited higher CAI values than the original yeast mRNAs, approaching the range of endogenous *E. coli* transcripts. This demonstrates that NUWA can effectively learn and apply host-specific codon usage patterns, generating mRNA sequences that are compositionally and structurally consistent with biological expression constraints.

### 2.4 NUWA shows potential to generate high-performance mRNA sequences

Developing an effective mRNA therapeutic often requires maximizing protein output per delivered mRNA molecule [28]. The translation efficiency, stability, and expression of mRNA are critical properties, especially in the context of mRNA vaccine applications, where they directly determine efficacy [29]. To explore NUWA’s capability in generating optimized mRNA sequences, we fine-tuned the pretrained model using translation efficiency(TE), stability, and expression datasets.

We first trained separate predictive models for TE, stability, and expression, which were subsequently used to curate a high-performance dataset by selecting sequences that exhibited consistently high scores in both ground-truth and model predictions (true positives). This curated dataset was then employed to fine-tune NUWA via the curriculum-masked language modeling (C-MLM) objective, thereby enhancing the model’s capacity to generate sequences with desired functional properties. Following fine-tuning, the enhanced model produced candidate mRNA sequences using the entropy-guided mRNA sequence generation pipeline (Figure 3.A). In the mRNA TE task, we employed the NUWA–bacteria model because the benchmark dataset is based on *E. coli*. In the mRNA stability task, we employed the NUWA–archaea model to leverage the unique mRNA coding patterns of archaeal species, which inhabit extreme environments and have evolved specialized mechanisms to maintain mRNA stability [30]. These characteristics may provide valuable insights for designing highly stable mRNA coding sequences.

**Fig. 3:**
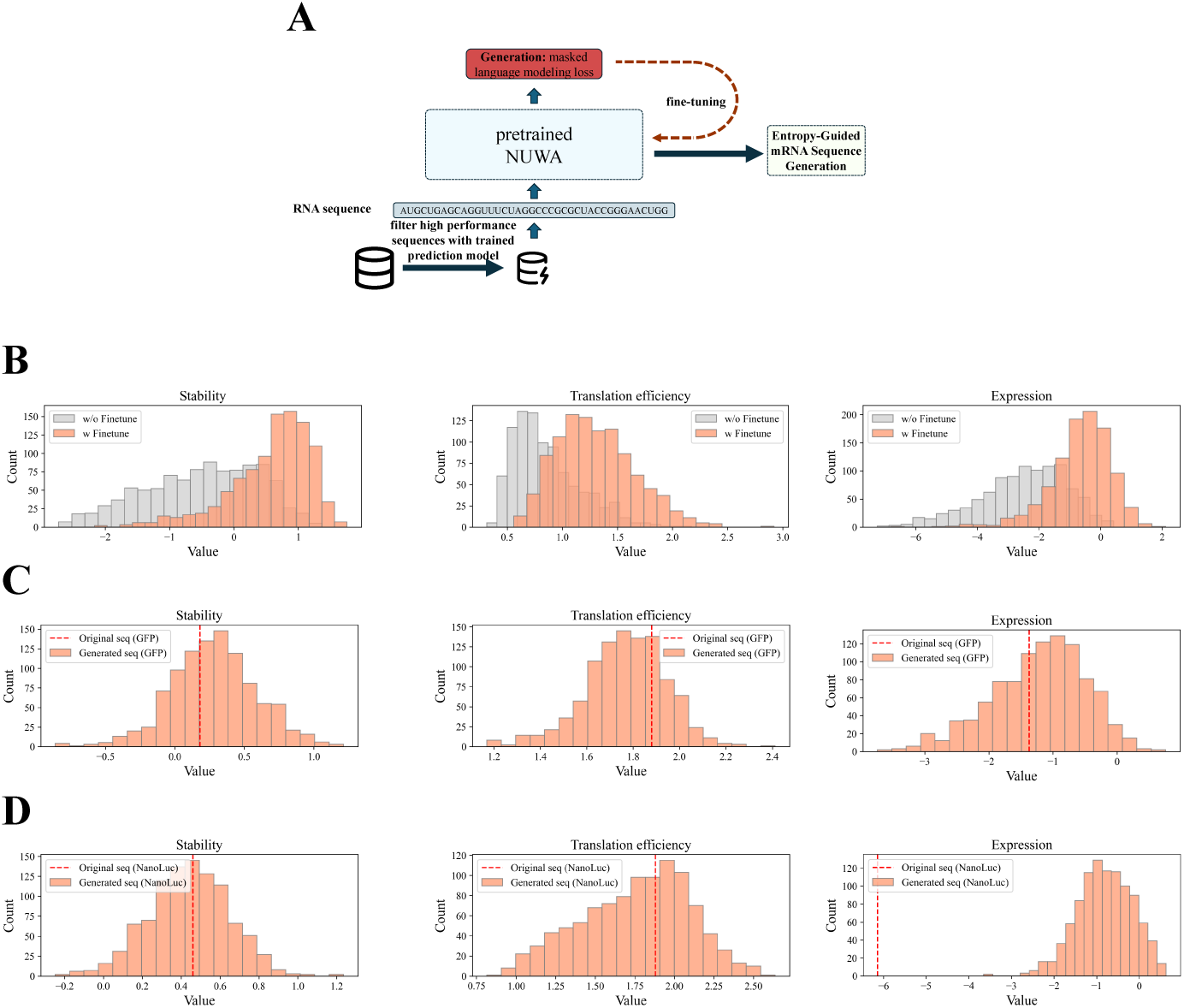
Fine-tuning NUWA for high-performance mRNA generation. (A) Schematic illustration of the fine-tuning process for the pretrained NUWA model, which was trained on experimentally validated, high-performance RNA datasets to enhance its generative capability. The fine-tuned model subsequently generates candidate mRNA sequences through an entropy-guided sampling pipeline. (B) Evaluation of mRNA stability, translation efficiency, and expression in generated sequences before and after fine-tuning, showing a rightward shift toward higher stability, translation efficiency, and expression values, respectively, after fine-tuning. (C) Distribution of predicted mRNA stability, translation efficiency, and expression generated by fine-tuned NUWA on the corresponding dataset. Here, the protein sequence was fixed to ensure that all mRNA sequences encode the GFP protein. The red dashed line indicates the predicted stability, translation efficiency, and expression value of the original GFP mRNA sequence. (D) Distribution of predicted mRNA stability, translation efficiency, and expression generated by fine-tuned NUWA on the corresponding dataset. Here, the protein sequence was fixed to ensure that all mRNA sequences encode the GFP protein. The red dashed line indicates the predicted stability, translation efficiency, and expression value of the original NanoLuc protein mRNA sequence, respectively.

As shown in Figure 3.B, the fine-tuned NUWA generated mRNAs with a distinct rightward shift in the stability-score, TE score, and expression distribution compared to the pretrained model, indicating improved intrinsic mRNA stability, TE, and expression, respectively. Comparative analysis revealed that the fine-tuned NUWA model generated mRNA sequences with significantly higher predicted stability, TE, and expression values compared to the pretrained model, respectively (Figure S5).

Furthermore, we evaluated the fine-tuned NUWA model on two representative reporter proteins, GFP and NanoLuc, which are commonly employed as biomarkers in molecular biology. The fine-tuned model successfully generated mRNA variants encoding these proteins with substantially higher predicted stability, translation efficiency (TE), and expression values than their respective original sequences Figure 3.C and Figure 3.D. This suggests that the fine-tuning process effectively improves both the performance and diversity of the generated mRNA sequence space.

We further extended this generation framework to applications in vaccines and therapeutics. Specifically, we selected three representative targets: the Spike protein mRNA sequence, the CD19 chimeric antigen receptor (CAR) mRNA sequence, and a degradation-promoting firefly luciferase variant mRNA sequence.

First, we evaluated recently published GEMORNA-designed mRNA sequences using our mRNA expression and stability predictor [26]. For all three proteins: firefly luciferase, Spike protein, and CD19 CAR, we observed that GEMORNA-designed mRNAs exhibited higher predicted expression and stability values (Figure S6), which is consistent with the claims in the GEMORNA paper that their model can generate mRNA sequences with enhanced translational capacity and stability [26].

Next, we used our fine-tuned NUWA to generate 1,000 mRNA sequences encoding each of the three proteins. For firefly luciferase, fine-tuned NUWA was able to generate sequences with higher predicted translation efficiency and expression than those designed by GEMORNA, classical, and commercial methods, although GEMORNAgenerated mRNAs also showed strong performance (Figure 4.A). For the CD19 CAR protein, fine-tuned NUWA was able to produce sequences with higher predicted stability and translation efficiency compared with GEMORNA and commercial benchmarks (Figure 4.B). For the Spike protein, fine-tuned NUWA was able to generate mRNA sequences with enhanced predicted stability, translation efficiency, and expression relative to GEMORNA and other publicly available commercial methods (Figure 4.C). These results illustrate that fine-tuned NUWA can generate biologically favorable mRNA variants and is capable of producing sequences that surpass established design baselines.

**Fig. 4:**
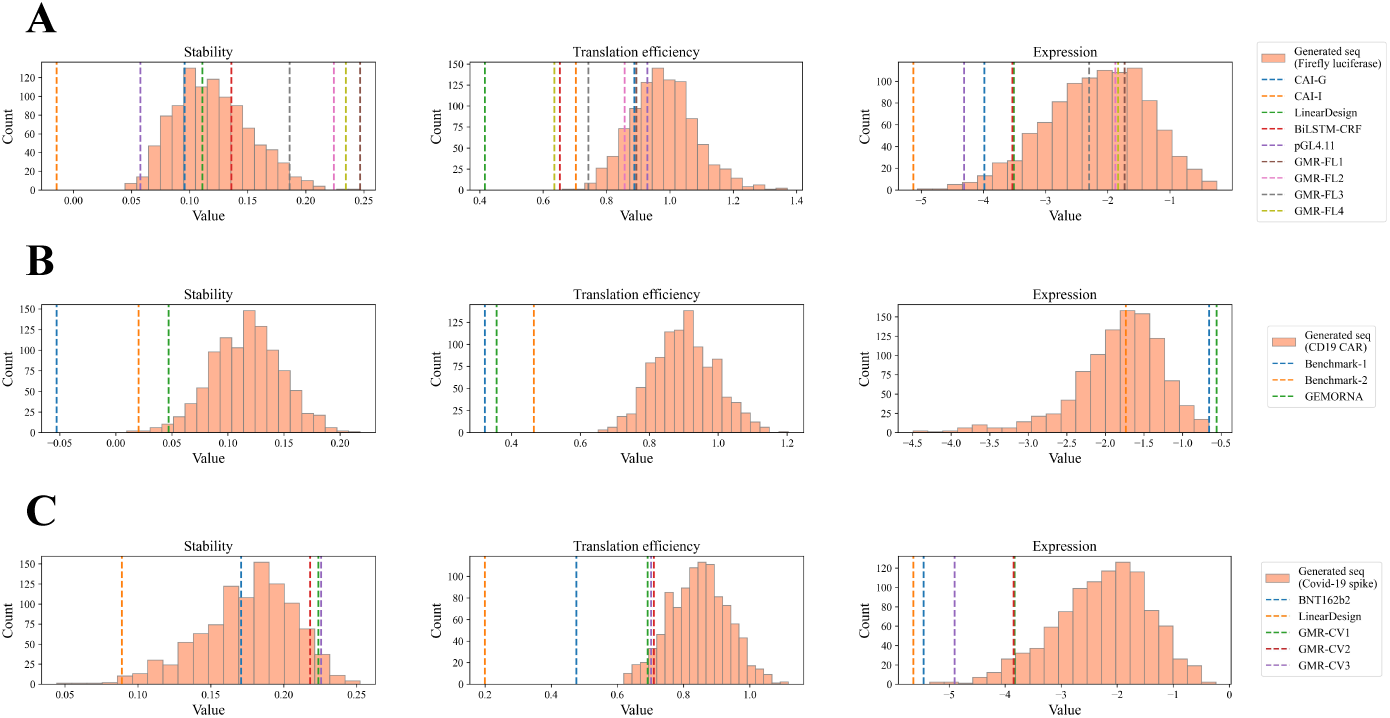
Predicted stability, translation efficiency, and expression of NUWA-generated mRNA sequences for three protein targets. Orange histograms represent the distribution of predicted values for NUWA-generated sequences. Vertical dashed lines indicate the corresponding values of published or benchmark mRNA designs, including GEMORNA-designed sequences and other publicly available or commercial design methods. Predicted stability, translation efficiency, and expression (left to right) of 1,000 mRNA sequences generated by fine-tuned NUWA for (A) firefly luciferase. (B) CD19 chimeric antigen receptor (CAR). (C) SARS-CoV-2 Spike protein.

Collectively, these results indicate that fine-tuning enables NUWA to internalize task-specific biological constraints and effectively generate mRNA sequences optimized for stability, translation efficiency, and expression, underscoring its potential as a tunable and powerful framework for de novo mRNA design in biomanufacturing, vaccine, and therapeutic applications.

### 2.5 NUWA shows strong capabilities in transfer learning across modalities for protein-related tasks

Building upon the mathematical principle that the codon-to–amino acid mapping is surjective but not injective, it is suggested that codon-based mRNA language models may capture more comprehensive biological information than protein language models [31]. This study extends a pretrained mRNA language model to protein-level tasks. This approach leverages the richer informational content embedded in codon sequences, which often exceeding that of amino acid representations, to enable cross-level modeling of protein structure and function.

Here, to further evaluate NUWA’s cross-modal generalization capacity, we explored its applicability to protein-related prediction tasks via transfer learning. Specifically, we repurposed the pretrained NUWA which originally trained on mRNA sequences by extracting its first encoder block and fine-tuning it on three protein-related datasets: fluorescence intensity prediction, protein stability prediction, and protein disorder prediction (Figure 5.A).

**Fig. 5:**
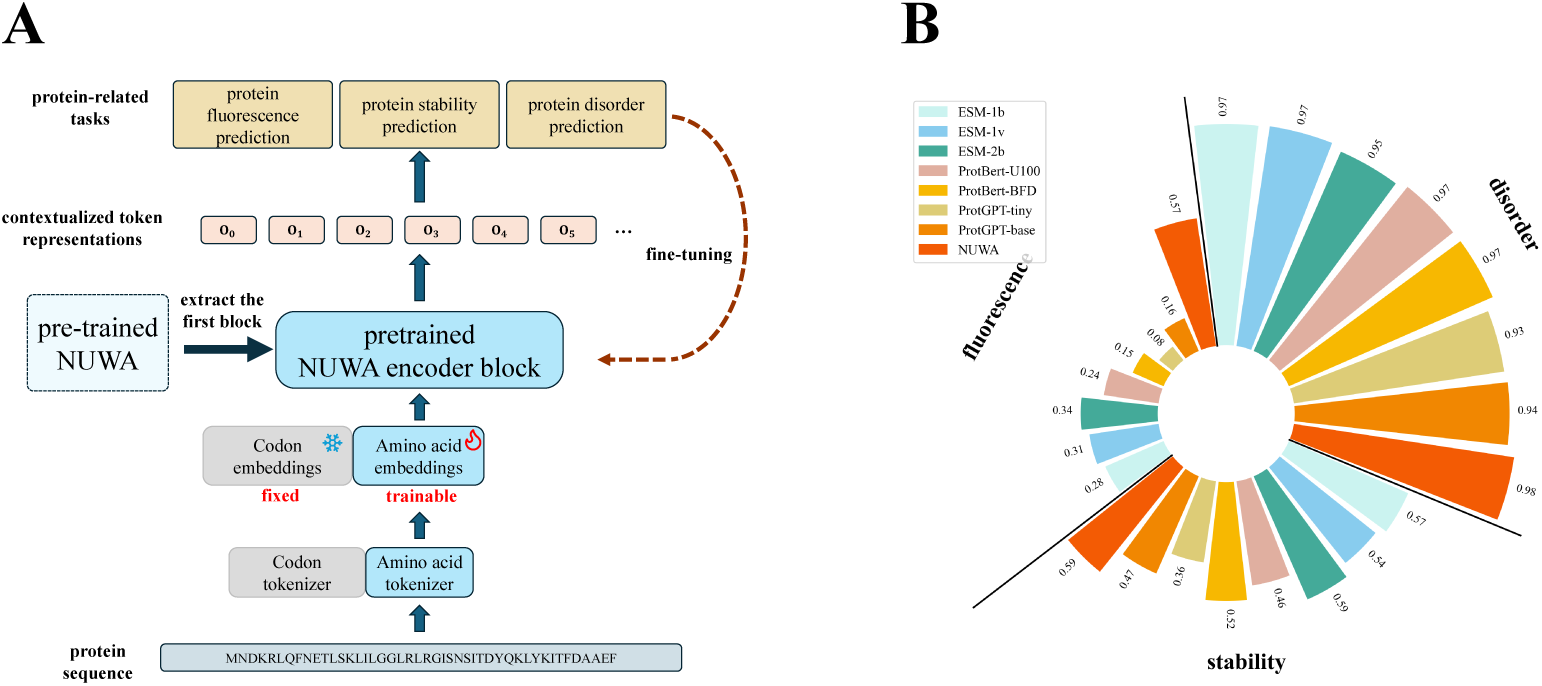
Cross-modal transfer learning of NUWA on protein-related tasks. (A) Schematic overview of applying NUWA for transfer learning in protein-level prediction tasks. The pretrained NUWA encoder—originally trained on large-scale mRNA coding sequences—was repurposed by extracting its first encoder block and fine-tuning it on protein datasets for three representative downstream tasks: fluorescence prediction, protein stability prediction, and protein disorder prediction. (B) Performance comparison of NUWA with state-of-the-art protein language models, including ESM-1b, ESM-1v, ESM-2b, ProtBert (BFD and U100), and ProtGPT (tiny and base), across the three benchmarks.

Remarkably, despite being trained exclusively on mRNA sequences, NUWA achieved comparable or superior performance to established protein language models such as ESM-1b, ESM-2b, ProtBert, and ProtGPT across all three benchmarks (Figure 5.B). In protein disorder prediction, NUWA reached a correlation score of 0.98, which matches specialized protein models. In stability prediction, NUWA maintained competitive performance (0.59) despite the domain shift from RNA to protein inputs. For fluorescence prediction, NUWA achieved a correlation of 0.57, again exceeding dedicated protein language models.

These results collectively highlight NUWA’s robust cross-modal representation ability, suggesting that codon-level contextual learning can capture biologically meaningful patterns transferable across molecular modalities, from mRNA to protein, without explicit retraining on protein corpora.

## 3 Discussion

mRNA occupies a central position in the central dogma, serving as the essential intermediary that transmits genetic information from DNA to the protein translation machinery [32]. Among biological applications of large language models (LLMs), protein language modeling has been the most extensively developed, largely due to its broad practical relevance in drug discovery, protein design, antibody engineering, and enzyme optimization, all of which are directly linked to the biopharmaceutical industry [33–36]. Although codons in mRNA sequences carry richer and more fine-grained biological information than amino acids in protein sequences, mRNA-focused language models remain relatively scarce [31]. In recent years, the rapid advancement of mRNA vaccines and RNA therapeutics has sparked growing interest and progress in mRNA language modeling [7, 24, 37].

In this study, we developed NUWA, the first large-scale mRNA language foundation model designed for unified sequence perception and generation. Our results demonstrate that NUWA not only captures the underlying statistical and biological principles encoded within mRNA sequences but also generalizes across diverse domains of life. By pretraining on over massive mRNA coding sequences spanning bacteria, archaea, and eukaryotes, NUWA effectively learned the evolutionary and codon usage patterns that define organism-specific translational rules. This broad taxonomic coverage enables the model to infer universal features of the mRNA language, while still preserving species-specific regulatory characteristics.

The fine-tuned versions of NUWA exhibited superior performance on a wide range of downstream perception tasks, including RNA structure prediction, function estimation, and engineering prediction. Notably, NUWA also exhibited strong transferability to cross-modal, protein-related tasks, suggesting that mRNA representations inherently encode information related to protein structure and function. This observation resonates with previous studies showing that codon usage is closely linked to protein folding, with experimental evidence demonstrating that synonymous codon substitutions can alter co-translational folding dynamics and ultimately affect the resulting protein’s conformation and function [15, 38–41]. This highlights the mRNA language model’s potential as a bridge between codon-level and protein-level understanding, paving the way for integrative biological modeling.

NUWA is the world’s first large language model to employ the BERT architecture for generation in the field of biology. NUWA demonstrates the ability to produce natural-like and biologically plausible mRNA sequences, reflecting its capacity to implicitly learn translation-relevant sequence features. When applied to heterologous protein expression, the model can further optimize the mRNA sequence encoding a given protein to match the codon usage bias of the target host species, thereby improving translational compatibility and efficiency. Unlike conventional codon optimization tools that rely on heuristic rules or predefined codon bias tables, NUWA achieves this through data-driven learning, highlighting its advantage as a knowledge-free yet biologically grounded generative framework for mRNA design.

Another important advantage of NUWA lies in its adaptability. The pretrained model can be efficiently fine-tuned on relatively small, task-specific datasets to generate functional mRNAs with desired properties, such as high translation efficiency, improved stability, or targeted expression profiles. In this study, our fine-tuned NUWA was able to generate mRNA sequences with high predicted translation efficiency, enhanced stability, and increased expression for five different protein targets, including GFP, NanoLuc, firefly luciferase, the SARS-CoV-2 Spike protein, and CD19 CAR. This flexibility allows researchers to rapidly prototype and explore mRNA designs tailored for synthetic biology, therapeutic development, and vaccine applications.

Overall, NUWA represents a conceptual and technical advancement in the field of biological language modeling. By unifying mRNA sequence perception and generation within a single framework, it provides a versatile foundation for programmable mRNA design. We anticipate that NUWA will not only facilitate a deeper understanding of RNA biology but also accelerate the development of next-generation mRNA-based therapeutics and synthetic constructs, marking a significant step toward AI-driven biological molecule design across all domains of life.

## 4 Methods

### 4.1 Datasets used for pretraining

The mRNA coding sequences used for pretraining NUWA were obtained from the NCBI RefSeq database (https://www.ncbi.nlm.nih.gov/datasets/genome/), which was accessed in August 2024. Specifically, the reference mRNA coding sequences of bacteria, archaea, and eukaryotes annotated by RefSeq were downloaded using the NCBI command-line tools (https://www.ncbi.nlm.nih.gov/datasets/docs/v2/command-line-tools/download-and-install/). In total, reference mRNA sequences from 19,676 bacterial, 4,688 eukaryotic, and 702 archaeal species were collected. For each domain, approximately 80 million bacterial, 2.1 million archaeal, and 83 million eukaryotic sequences were retrieved, respectively.

### 4.2 Datasets used for RNA-related downstream tasks

Tables 2 and 4 summarize the datasets for the tasks used to compare NUWA with CodonBERT and BEACON, respectively. These tasks provide a broad assessment covering three major categories. These range from structural prediction, which focuses on inferring the two or three dimensional architecture of the RNA molecule, to functional prediction, which aims to predict functional outcomes given the input mRNA sequences, and finally, engineering prediction, which assesses the predictability of designed biological systems. Specifically, the benchmarking tasks detailed in Table 2 cover functional prediction, and engineering prediction:

- Functional Prediction:

– **mRFP Expression** [42] quantifies protein production levels in *Escherichia coli* for a library of gene variants.
– **Fungal Expression** [43] contains coding sequences, each greater than 150 base pairs (bp), curated from a diverse set of fungal genomes.
– ***E. coli* Protein Expression** [44] includes experimental measurements of protein abundance in *E. coli*, with corresponding mRNA sequences categorized into three discrete expression levels: low (2,308 sequences), medium (2,067 sequences), and high (1,973 sequences).
– **mRNA Stability** [45] comprises empirical mRNA half-life measurements across multiple model organisms, including human, mouse, frog, and fish species.
– **SARS-CoV-2 Vaccine Degradation** [46] consists of chemically optimized mRNA sequences targeting structural features, stability, and translational efficiency. The prediction target for this dataset is the average deg Mg 50C value per nucleotide.
- Engineering Prediction:

– **Tc-Riboswitch** [47] features tetracycline (Tc) riboswitch dimer sequences engineered upstream of a green fluorescent protein (GFP) mRNA reporter.
The benchmarking tasks detailed in Table 4 cover structural prediction, functional prediction, and engineering prediction:
- Structural Prediction:

– **Secondary Structure Prediction (SSP)** [48] identifies base-paired and unpaired regions within the RNA sequence.
– **Contact Map Prediction (CMP)** [49] identifies nucleotide pairs that are in close spatial proximity in the folded structure.
– **Distance Map Prediction (DMP)** [49] estimates the physical distance between all nucleotide pairs.
- Functional Prediction:

– **APA Isoform Prediction (APA)** [50] predicts the usage ratio of proximal polyadenylation (poly(A)) sites.
– **Non-coding RNA Function Classification (ncRNA)** [51, 52] categorizes ncRNAs into functional types, such as microRNAs (miRNAs), long non-coding RNAs (lncRNAs), and small interfering RNAs (siRNAs).
– **Modification Prediction (Modif)** [53] predicts the occurrence of twelve common RNA chemical modifications from a given RNA sequence.
– **Mean Ribosome Loading (MRL)** [54] predicts the Mean Ribosome Loading value for input mRNA sequences, a proxy for translational efficiency.
- Engineering Prediction:

– **Programmable RNA Switches (PRS)** [55] predicts the activity states of synthetic RNA regulatory switches.
– **CRISPR On-Target Prediction (CRI-On)** [56] evaluates the on-target cleavage efficiency of single-guide RNAs (sgRNAs).
– **CRISPR Off-Target Prediction (CRI-Off)** [56] assesses the likelihood of CRISPR-induced mutations occurring at unintended genomic locations.

**Table 4:**
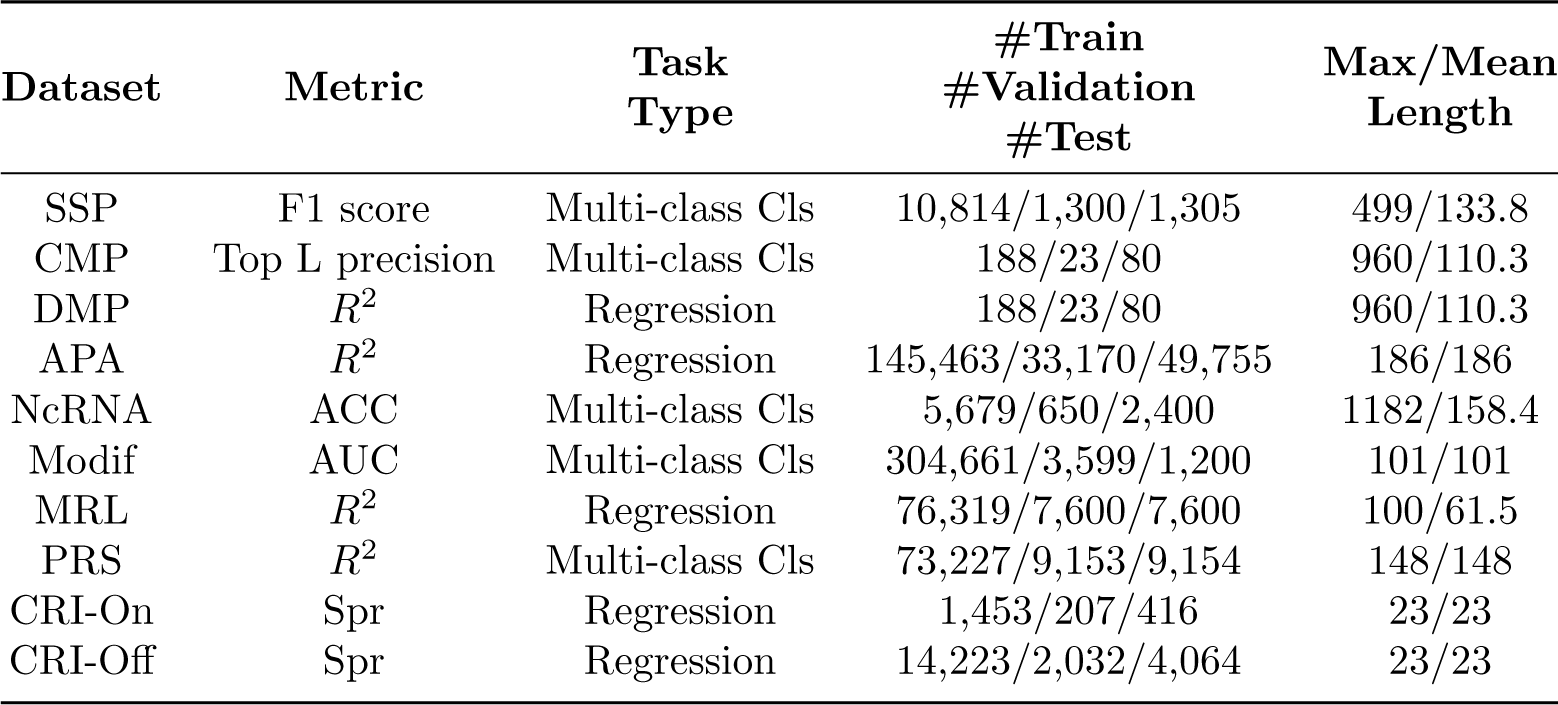
Overview of task datasets used in Table 3. “Cls” indicates “classification”, “Spr” indicates spearman correlation coefficient, “ACC” indicates accuracy, and “R^2^” indicate coefficient of determination.

| Dataset | Metric | Task Type | #Train<br>#Validation<br>#Test | Max/Mean Length |
| --- | --- | --- | --- | --- |
| SSP | F1 score | Multi-class Cls | 10,814/1,300/1,305 | 499/133.8 |
| CMP | Top L precision | Multi-class Cls | 188/23/80 | 960/110.3 |
| DMP | $R^2$ | Regression | 188/23/80 | 960/110.3 |
| APA | $R^2$ | Regression | 145,463/33,170/49,755 | 186/186 |
| NcRNA | ACC | Multi-class Cls | 5,679/650/2,400 | 1182/158.4 |
| Modif | AUC | Multi-class Cls | 304,661/3,599/1,200 | 101/101 |
| MRL | $R^2$ | Regression | 76,319/7,600/7,600 | 100/61.5 |
| PRS | $R^2$ | Multi-class Cls | 73,227/9,153/9,154 | 148/148 |
| CRI-On | Spr | Regression | 1,453/207/416 | 23/23 |
| CRI-Off | Spr | Regression | 14,223/2,032/4,064 | 23/23 |

### 4.3 Datasets used for transfer learning in protein-related tasks

Table 5 provides the dataset summary of protein-related tasks:

- **Protein fluorescence** [57] predicts the fluorescence level of *Aequorea victoria* Green Fluorescent Protein (avGFP) variants.
- **Protein stability** [58, 59] aims to predict the stability of a protein, a crucial measure of its ability to maintain structural and functional integrity.
- **Protein disorder** [60] involves the identification of intrinsically disordered regions.

**Table 5:** Dataset overview of protein-related tasks used in Figure 5.A. “Cls” indicates “classification”, “Spr” indicates spearman correlation coefficient, and “ACC” indicates accuracy.

| Dataset | Metric | Task Type | #Train<br>#Validation<br>#Test | Max/Mean Length |
| --- | --- | --- | --- | --- |
| Protein fluorescence | Spr | Regression | 21,446/5,362/27,217 | 237/237 |
| Protein stability | Spr | Regression | 53,614/2,512/12,851 | 50/45 |
| Protein disorder | ACC | Binary Cls | 8,678/2,170/513 | 1632/257.3 |

### 4.4 Protein sequence and mRNA coding sequence used for generation and evaluation

For mRNA coding sequence generation for target proteins, the GFP and NanoLuc protein sequences and their corresponding wild-type mRNA sequences were downloaded from NCBI. Designed mRNA coding sequences for firefly luciferase, the SARS-CoV-2 Spike protein, and the CD19 CAR, which were produced by GEMORNA and other publicly available commercial design methods, and evaluated using our stability, translation efficiency, and expression predictors, which were obtained from [26]. The corresponding protein sequences were translated using our in-house translation algorithm.

### 4.5 Details of NUWA

#### Input representation

Our model operates based on codon-level tokenization. A codon is defined as a triplet of three adjacent nucleotides. For each position within a codon, there are four nucleotide options: Adenine (A), Uracil (U), Guanine (G), and Cytosine (C). This results in a total of 4^3^ = 64 possible codon combinations, constituting the standard genetic code. Moreover, four special tokens are incorporated to facilitate model training and downstream tasks: a classifier token ([`CLS`]), a separator token ([`SEP`]), a padding token ([`PAD`]), and a masking token ([`MASK`]). Consequently, the final vocabulary of our language model comprises 68 unique tokens.

#### Model architecture

As illustrated in Figure 1, our NUWA framework accepts a batch of mRNA sequences as input. For a given mRNA sequence belonging to the i-th species, the sequence is initially segmented into a series of codons. These codons are then tokenized to form the input sequence for the BERT model. The sequence construction follows standard practices: a [`CLS`] token is prepended to the start to enable the aggregation of sequence-level representations, and a [`SEP`] token is appended to the end to demarcate the sequence boundary. Additionally, sequences are padded with the [`PAD`] token to ensure fixed-length batch processing. Thereby these practices result in learnable codon embeddings. To encode the sequential order, learnable positional embeddings are added to the token embeddings. Furthermore, a unique, learnable segment embedding corresponding to the i-th species is added to all tokens in the input, providing crucial class information to the model. In summary, the model’s final input embedding is constructed by summing three distinct components: the codon embedding, the positional embedding, and the segment embedding of i-th species.

The combined input embedding is subsequently processed by the BERT model, which comprises a stack of 12 identical transformer encoder blocks, as detailed in Figure 1.B. Each encoder layer utilizes 12 multi-head self-attention (MHSA) layers to compute comprehensive, bidirectional representations of the input sequence, operating with a hidden state dimension of 768. MHSA dynamically weighs and integrates contextual information from all tokens across the entire input sequence, allowing the model to capture both local and long-range interactions. Following the multi-head selfattention mechanisms, a feed-forward network (FFN) is applied to refine the extracted feature representations. Throughout the architecture, residual connections and layer normalization are consistently employed within both the MHSA and FFN to facilitate feature flow. After processing through the entire encoder stack, the model outputs contextualized token representations. During the pretraining phase, these representations are utilized for objectives such as predicting masked tokens via a softmax classifier and performing other self-supervised objectives.

#### Supervised contrastive loss (SCL)

To equip NUWA with robust discriminative perception and enhanced generation capabilities, we introduce two primary innovations during the pretraining phase: the supervised contrastive loss (SCL) and a curriculum masked language modeling (C-MLM) objective. Specifically, the SCL is designed to enforce the learning of highly discriminative and semantically meaningful feature representations, thereby enhancing the discriminative perception capacity of the model. It achieves this by pulling the feature embeddings of samples belonging to the same class closer together in the representation space, while simultaneously pushing those from different classes further apart. Let **z***_i_* ∈ ℝ*^d^* denote the ℓ_2_-normalized [`CLS`] feature embedding of the i-th sample within a training batch, and let *y_i_* be its corresponding class label. The similarity between any two embeddings is calculated using the cosine similarity function, scaled by a temperature parameter *τ*. A positive pair is defined as any two samples sharing the identical class label. The SCL objective is to maximize the similarity of these positive pairs relative to all other samples in the batch. Formally, the SCL loss for a batch of *N* samples is defined as:

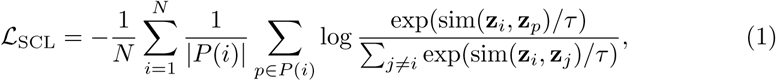

where *P*(*i*) denotes the set of indices of all samples in the batch that share the same class label as sample *i* (i.e., *y_j_* = *y_i_*), excluding the anchor sample *i* itself. The sim(·, ·) denotes the cosine similarity function, and the exp(·) indicates the exponential function. This objective explicitly promotes intra-class compactness and inter-class separability within the learned representation space to enhance the quality of the contextualized embeddings. To ensure the effective computation of the SCL, which requires a balanced and diverse set of positive and negative pairs, we enforce a specific batch sampling constraint. In each batch, we randomly sample *K* = 64 distinct classes and ensure that each selected class has 4 samples.

#### Curriculum masked language modeling (C-MLM)

In complement to the SCL, we also adopt a curriculum masked language modeling (C-MLM) objective to specifically enhance the model’s generative capabilities, which incorporates a curriculum learning strategy to progressively increase the difficulty of the masked language modeling (MLM) objective, thereby incrementally improving the sequence generation capacity. Building on the MLM, C-MLM operates by randomly selecting a subset of tokens in the input sequence and replacing them with the masking token ([`MASK`]). The model is then trained to predict the original tokens at these masked positions, leveraging the contextual information provided by the unmasked tokens. For each masked position, the model outputs a probability distribution over the entire vocabulary. The masked language modeling loss, denoted 𝓛_MLM_*_r_*, is calculated as the average cross-entropy loss over all masked tokens:

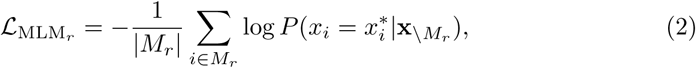

where *M_r_* is the set of indices corresponding to the *r*% masked positions, 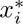 is the true original token at position *i*, and **x**_\*M*_*_r_* represents the set of non-masked tokens providing the context. For gradually increasing the learning difficulty, the masking probability *r* is dynamically adjusted at the end of each training step according to a linear scheduling function:

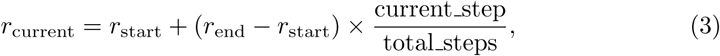

where *r*_start_ = 0.15 and *r*_end_ = 0.5 are the initial and final masking probabilities, respectively. current step and total steps track the current training iteration and the total number of iterations. At the beginning of training, the low initial masking probability (e.g., 0.15) presents an easier token prediction task, allowing the model to quickly establish reliable sequence semantics from an abundance of unmasked tokens. This setting biases the model towards developing a highly discriminative understanding of local and global sequence features. Conversely, as training progresses, the linear increase in *r* (e.g., 0.5) introduces a more challenging token prediction task progressively. A higher masking ratio necessitates greater reliance on sparse contextual clues, forcing the model to infer a larger proportion of missing content. This setting biases the model to move beyond reliance on local cues and develop a deeper generative understanding of the entire sequence structure to accurately predict the missing information, thereby significantly enhancing its overall generative capacity.

### 4.6 Details of pretraining process

For the pretraining phase, the AdamW optimizer [61] was used with an initial learning rate of 1 × 10^−5^, which was adjusted using a linear decay scheduler, a weight decay of 1 × 10^−2^, and 1,000 warmup steps. The archaea and bacteria models were pretrained on all collected mRNA sequences. Given the massive scale of the eukaryote data, we employed a random sampling strategy, selecting 40% of the data from each species to form the final pretraining set. In each batch, we randomly sample 64 distinct classes, with 4 samples per class, yielding a final per-GPU batch size of 256. Sequence length limits were set to 1,024 codons for the archaea and bacteria models, and 512 codons for the eukaryote model. Sequences exceeding these limits were truncated. All training was implemented within the PyTorch framework. The pretrained model encompasses approximately 110 million parameters.

### 4.7 Implementation details of fine-tuning process for perception task

Comparison with CodonBERT. For the tasks summarized in Table 1, all models were optimized using the AdamW optimizer (with initial learning rates of 5×10^−5^, *β*_1_ = 0.9, *β*_2_ = 0.999, weight decay of 0.01), coupled with a linear decay learning rate scheduler. Training was limited to a maximum of 1,000 steps, with a batch size of 8. The choice of pretrained model was matched to the biological context of the downstream tasks: the NUWA-bacteria model is used for mRFP expression, *E. coli* protein expression, Tc-riboswitch, and SARS-CoV-2 vaccine degradation tasks. Conversely, the NUWA-eukaryote model is applied to the Fungal expression and mRNA stability tasks. During the fine-tuning process, if the species of the dataset is known, we assign explicit class labels accordingly. Otherwise, we designate the first class as a virtual default label. Thanks to the strong discriminative capability introduced by SCL, we observe that even such a defective value can still lead to satisfactory performance.

Comparison with BEACON. To ensure a fair and consistent comparison for the results presented in Table 3, we adopted the fine-tuning setup reported in [62]. All models were optimized using the AdamW optimizer (*β*_1_ = 0.9, *β*_2_ = 0.999, weight decay of 0.01), with a linear decay learning rate scheduler. The fine-tuning hyperparameter settings were tailored for each dataset. The structural prediction models SSP, CMP, and DMP were trained for 100 epochs, with initial learning rates of 3 × 10^−5^, 3 × 10^−7^, and 5 × 10^−5^, respectively. Effective batch sizes varied significantly: SSP uses a batch size of 4 with a gradient accumulation of 4; CMP uses a batch size of 1 with accumulation of 8; and DMP uses a batch size of 1 with accumulation of 2. Functional and engineering tasks, including APA, NcRNA, Modif, MRL, PRS, CRIOn, and CRI-Off, were trained for 30 epochs. The initial learning rate was set to 5 × 10^−5^ for APA, NcRNA, and MRL, and 3 × 10^−5^ for Modif, while PRS, CRI-On, and CRI-Off use 1 × 10^−5^. These tasks employed batch sizes from 16 to 128 (APA: 32, NcRNA: 16, Modif: 32, MRL: 128, PRS: 32, CRI-On: 16, CRI-Off: 16) with gradient accumulation of 2. Furthermore, we maintained the domain-specific matching between the pretrained model and the downstream tasks. The NUWA–bacteria model is used for SSP, CMP, DMP, ncRNA, Modif, CRI-On, and CRI-Off. The NUWA–eukaryote model is adopted for APA, MRL, PRS, CRI-On, and CRI-Off.

For the fine-tuning process for transfer learning in protein tasks, we adopted the fine-tuning setup from [63] to ensure a fair comparative analysis. Specifically, the model was optimized using AdamW (β_1_ = 0.9, β_2_ = 0.999, weight decay 0.01) with an initial learning rate of 6×10^−4^, a batch size of 64, and a total of 100 epochs, utilizing a cosine decay learning rate scheduler. As shown in Figure 5.A, we utilized the first encoder block of the pretrained NUWA model as the backbone for these transfer learning tasks. To accommodate the protein sequences, we set a new amino acid tokenizer (20 amino acid tokens + 4 special tokens), which results new trainable amino acid embeddings. Here, we adopt the NUWA-archaea model. This choice is based on the principle that shallow layers generally learn to extract fundamental, highly transferable features, such as basic sequence structure and local dependencies [64, 65], common to both nucleic acids and proteins. While the deeper layers encode mRNA-specific semantic representations [66], leading to a mismatch when applied to the protein domain.

### 4.8 The generation algorithm of NUWA

#### Algorithm 1 Entropy-Guided mRNA Sequence Generation

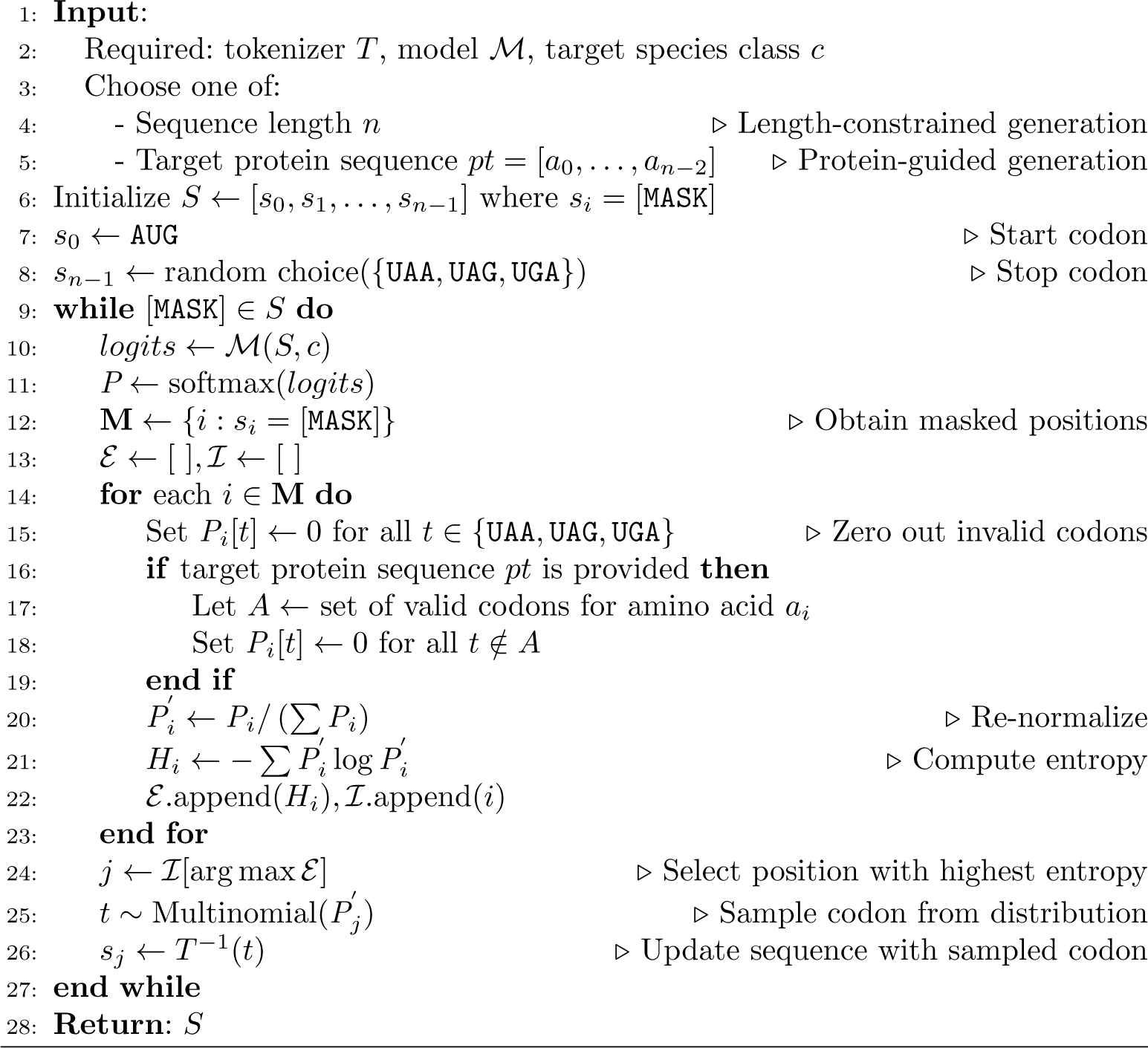

In this work, we propose a unified entropy-guided mRNA sequence generation approach, which supports both length-constrained and protein-guided generation strategies. The generation process is driven by a pretrained BERT-like model ℳ, and incorporates biological constraints to ensure the validity of the synthesized mRNA sequences.

Algorithm 1 outlines the entropy-guided generation procedure. For length-constrained mRNA generation, the algorithm takes as input a desired sequence length n, a tokenizer *T*, a model ℳ, and a target class label c. Additionally, for protein-guided generation, instead of length *n*, a target protein sequence *pt* = [a_0_, a_2_, · · ·, a*_n_*_−2_] is provided, where each *a_i_* is an amino acid corresponding to the codon position *i* in the mRNA sequence.

The sequence *S* = [*s*_0_, *s*_1_, …, *s_n_*_−1_] is initialized with all positions masked. The first codon at position s_0_ is set to `AUG`, and the last codon at position s*_n_*_−1_ is randomly assigned a valid stop codon from the set {UAA, UAG, UGA}, following biological constraints. The generation proceeds iteratively. At each step, the model ℳ predicts token logits for all positions in *S*, conditioned on the current sequence and class *c*. These logits are converted into probability distributions **P** via the softmax function. For each masked position i, the corresponding distribution **P***_i_* is modified by applying biological constraints:

- In length-constrained generation, stop codons are removed from the internal positions by zeroing out their probabilities.
- In protein-guided generation, in addition to removing stop codons, codons that do not translate to the target amino acid a*_i_*_−1_ are also masked out. For example, for amino acid `Phenylalanine`, all codons except for `UUU` and `UUC` will be zeroed out.

The constrained distribution **P***_i_* is then re-normalized to ensure it forms a valid probability distribution 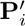, from which the entropy H*_i_* is computed. Entropy serves as a measure of the model’s uncertainty at each position. Among all masked positions, the one with the highest entropy is selected, and a codon is sampled from its normalized distribution 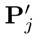. This entropy-guided mRNA sequence generation approach intentionally selects the position with the highest entropy to maximize diversity and offer the greatest flexibility during the generation process. The sampled codon is then inserted into the sequence at position j. The process repeats until all positions in S are filled. The final output is a biologically valid mRNA sequence S that begins with a start codon, ends with a valid stop codon, and satisfies the specified constraints. In length-constrained mode, the sequence conforms to the given length and class label. In protein-guided mode, the generated mRNA translates exactly into the target protein sequence *pt*, while also adhering to the class condition.

### 4.9 Implementation details of generating high-performance mRNA sequences

To enable the NUWA models to generate high-performance mRNA sequences, we implemented a specialized procedure involving data curation and targeted fine-tuning. First, to create a high-quality dataset for training the NUWA model to generate high-performance mRNA sequences, we required accurate models to identify high-performing sequences. To this end, we trained separate predictive models for mRNA stability, translation efficiency, and expression. These predictive models were trained by employing the same fine-tuning settings as described in Table 1. The dataset for the stability model followed the configuration outlined in Table 2. The TE dataset [67] was partitioned sequentially into 70% training, 15% validation, and 15% test sets. Then, utilizing the predictive models, we curated a specialized dataset composed exclusively of high-performing sequences. The selection criterion was to identify “true positives”, i.e., sequences that exhibited high values in both the ground-truth data and the model predictions. For stability, we retain the sequence with both the true stability value and the predicted value to be greater than 0.5. While for TE, we retain the sequence if its true TE value exceeds 1 and its predicted TE surpasses 1. The expression adopted the same setting with stability. Subsequently, this curated set of high-performance sequences was employed to fine-tune the pretrained NUWA model via C-MLM. This step specifically enhances the model’s capacity to generate sequences with these desired high-performance characteristics. For this fine-tuning process, we adopted the setting of the original pretraining process, except for running for 200 total training epochs. Finally, the optimized NUWA model was employed within an entropy-guided mRNA generation pipeline to generate high-performance target mRNA sequences. Here, we conduct both length-constrained generation and protein-guided generation. For the length-constrained approach, we addressed three tasks: in the mRNA TE task, we employed the NUWA–bacteria model, setting the species to *E. coli* because the benchmark dataset is based on *E. coli*; in the mRNA stability task, we used the NUWA–archaea model, selecting the radiation-resistant organism *Thermococcus gammatolerans*. For the protein-guided approach, we specified the protein sequences for green fluorescent protein (GFP), NanoLuc, firefly luciferase, spike protein, and CD19 CAR. The generated mRNA sequences encoding these target proteins were subsequently evaluated for stability, translation efficiency, and expression prediction.

## Acknowledgements

- This work was partly supported by JST CREST JPMJCR23N1. The computations were partially performed on the NIG supercomputer at ROIS National Institute of Genetics, HOKUSHIN supercomputer at the Department of Data Science, Kitasato University.
- Conflict of interest/Competing interests The authors declare no competing interests.
- Data availability. All data used to train the model are available at https://www.ncbi.nlm.nih.gov/datasets/genome/. Data for Table 1 are available at https://github.com/Sanofi-Public/CodonBERT. Data for Table 3 are available at https://github.com/terry-r123/RNABenchmark.
- Code availability. The code is available in https://github.com/zysxmu/NUWA.

## Appendix A

### Visualization of projected embedding

Figure S1 presents the visualization of projected feature embedding from NUWA-archaea, NUWA-bacteria, and NUWA-eukaryote.

**Fig. S1:**
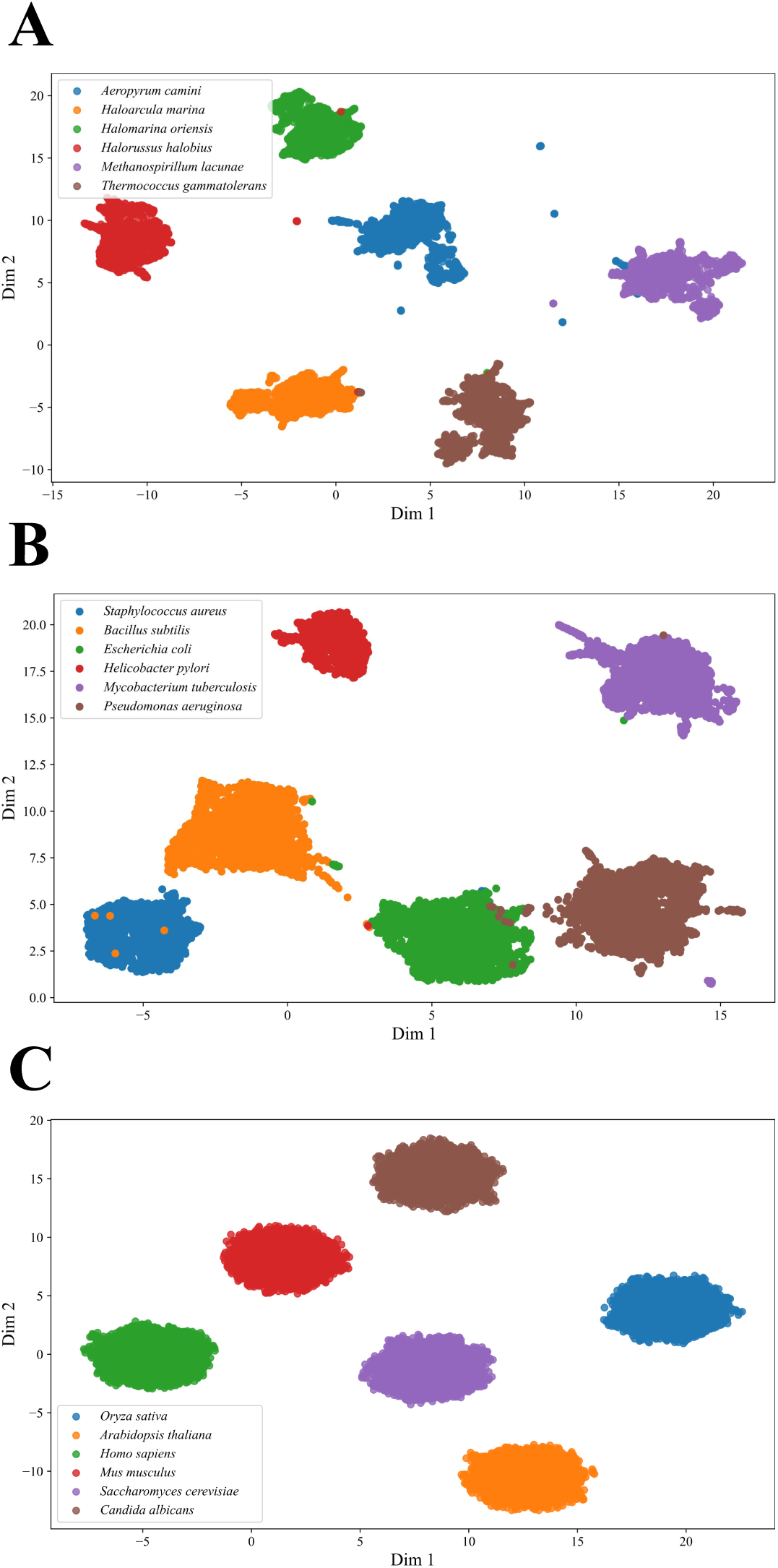
UMAP [68] visualization of projected feature embedding. (A) NUWA-archaea. (B) NUWA-bacteria. (C) NUWA-eukaryote.

## Appendix B

### Train and Evaluation Loss

Figure S2 presents the pretraining loss of NUWA-archaea, NUWA-bacteria, and NUWA-eukaryote.

## Appendix C

### Correlation plots of mRNA stability, translation efficiency, and expression tasks

Figure S3 illustrates the correlation results on the test set for the mRNA stability, translation efficiency, and expression tasks.

## Appendix D

### Distribution of mRNA stability and translation efficiency, and expression datasets.

Figure S4 illustrates the distribution of mRNA stability and translation efficiency, and expression datasets.

## Appendix E

### Distribution of Generated high-performance mRNA sequences

Figure S5 illustrates the distribution of evaluation results of generated mRNA in the mRNA stability, translation efficiency, and expression tasks. This figure serves as a supplementary analysis to Figure 3.B, Figure 3.C, and Figure 3.D, providing box plots along with corresponding p-values.

## Appendix F

### Stability and Expression Prediction Results for mRNA Coding Sequences

Figure S6 illustrates the evaluation results of mRNA coding sequence encoding for firefly luciferase (A), Spike protein (B), and CD19 CAR (C) by using our stability and expression predictor.

**Fig. S2:**
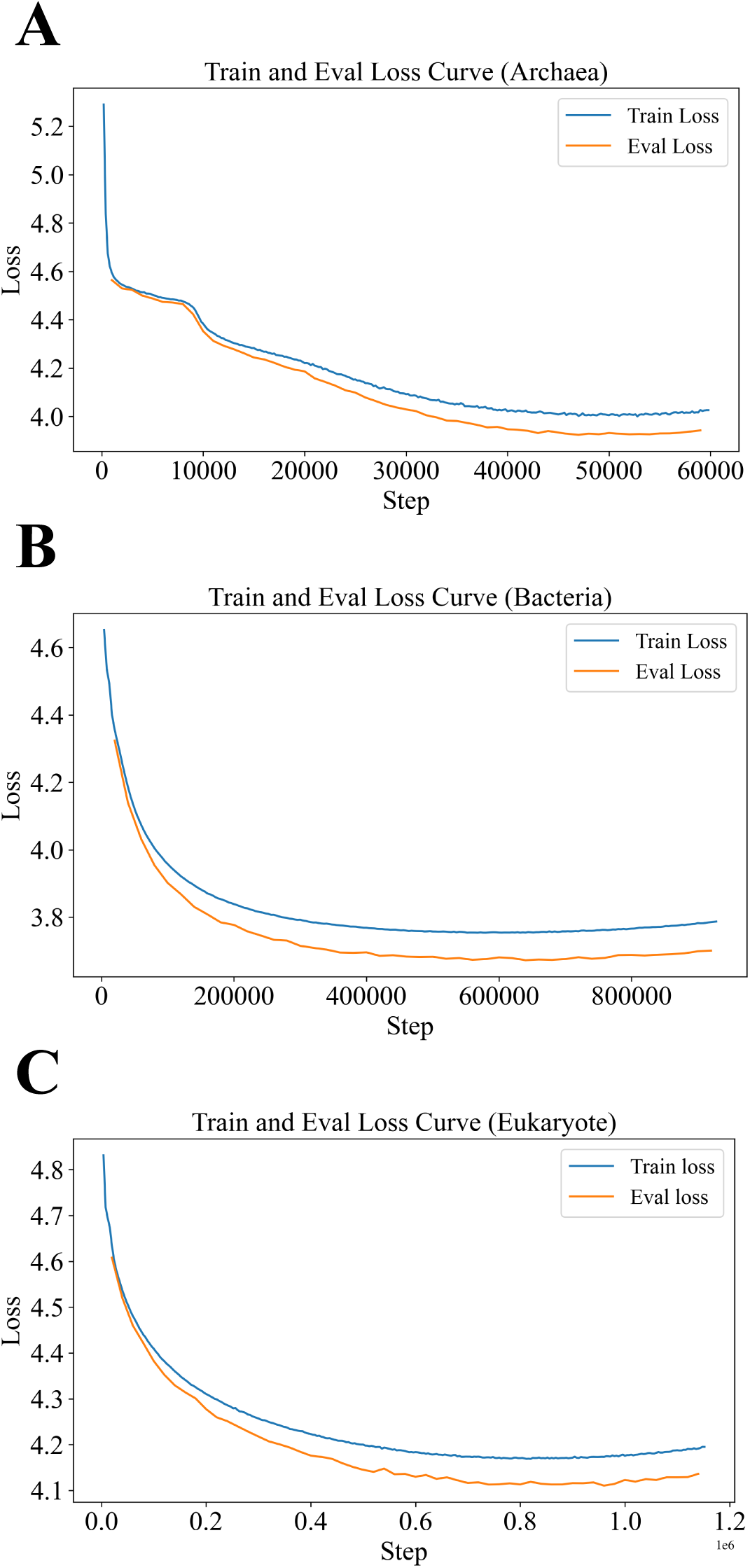
Pretraining loss. (A) NUWA-archaea. (B) NUWA-bacteria. (C) NUWAeukaryote.

**Fig. S3:**
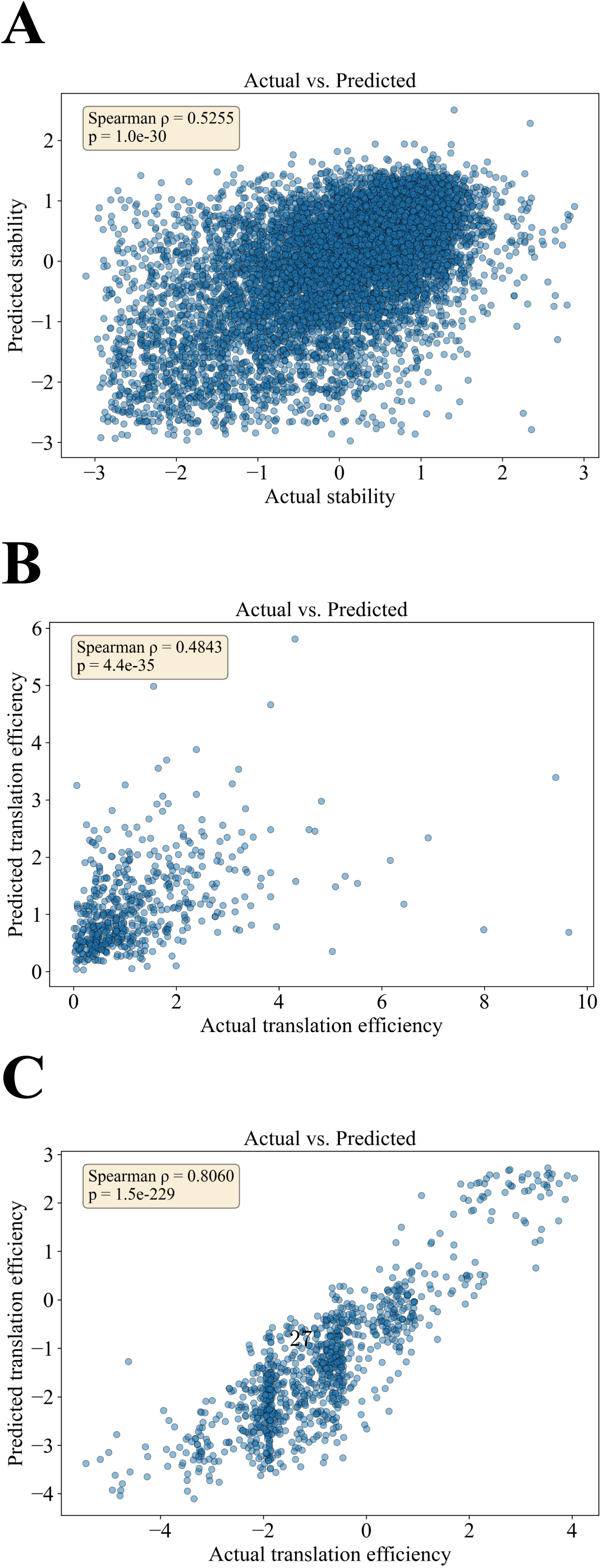
Evaluation results on test set. (A) mRNA stability; (B) mRAN translation efficiency (TE); (C) mRNA expression.

**Fig. S4:**
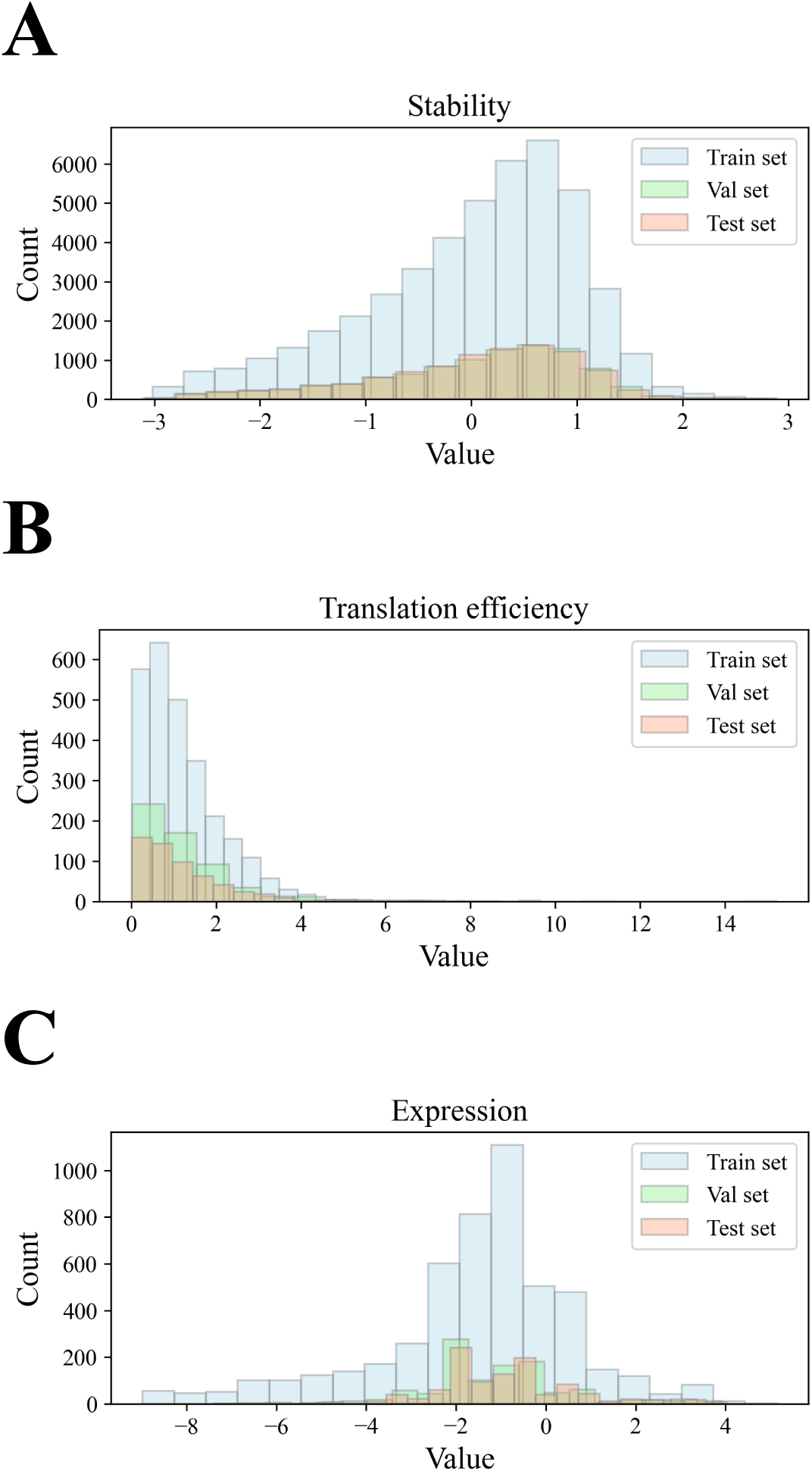
Distribution of mRNA stability and translation efficiency datasets. (A) mRNA stability; (B) mRNA translation efficiency (TE); (C) mRNA expression.

**Fig. S5:**
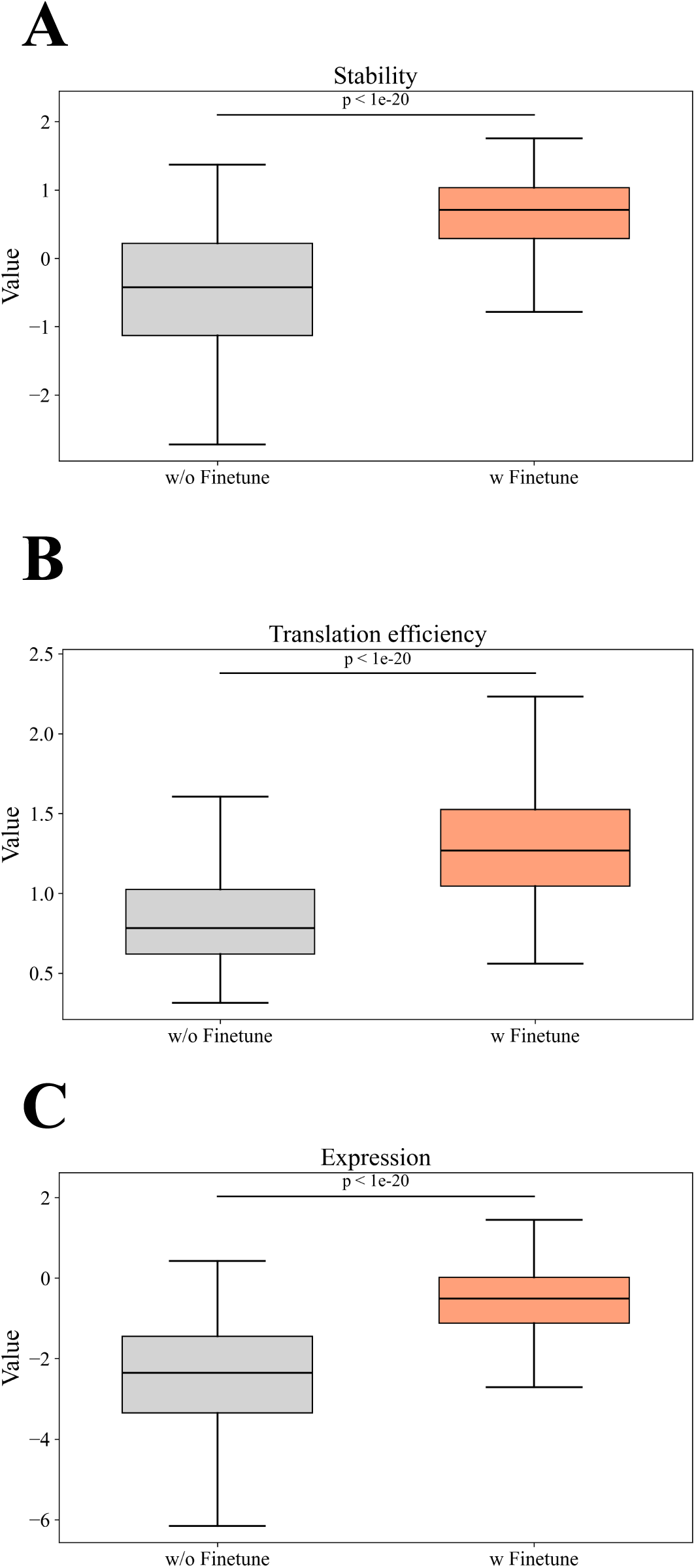
Distribution of generated mRNA on mRNA stability and translation efficiency tasks. (A) mRNA stability; (B) mRNA translation efficiency (TE); (C) mRNA expression.

**Fig. S6:**
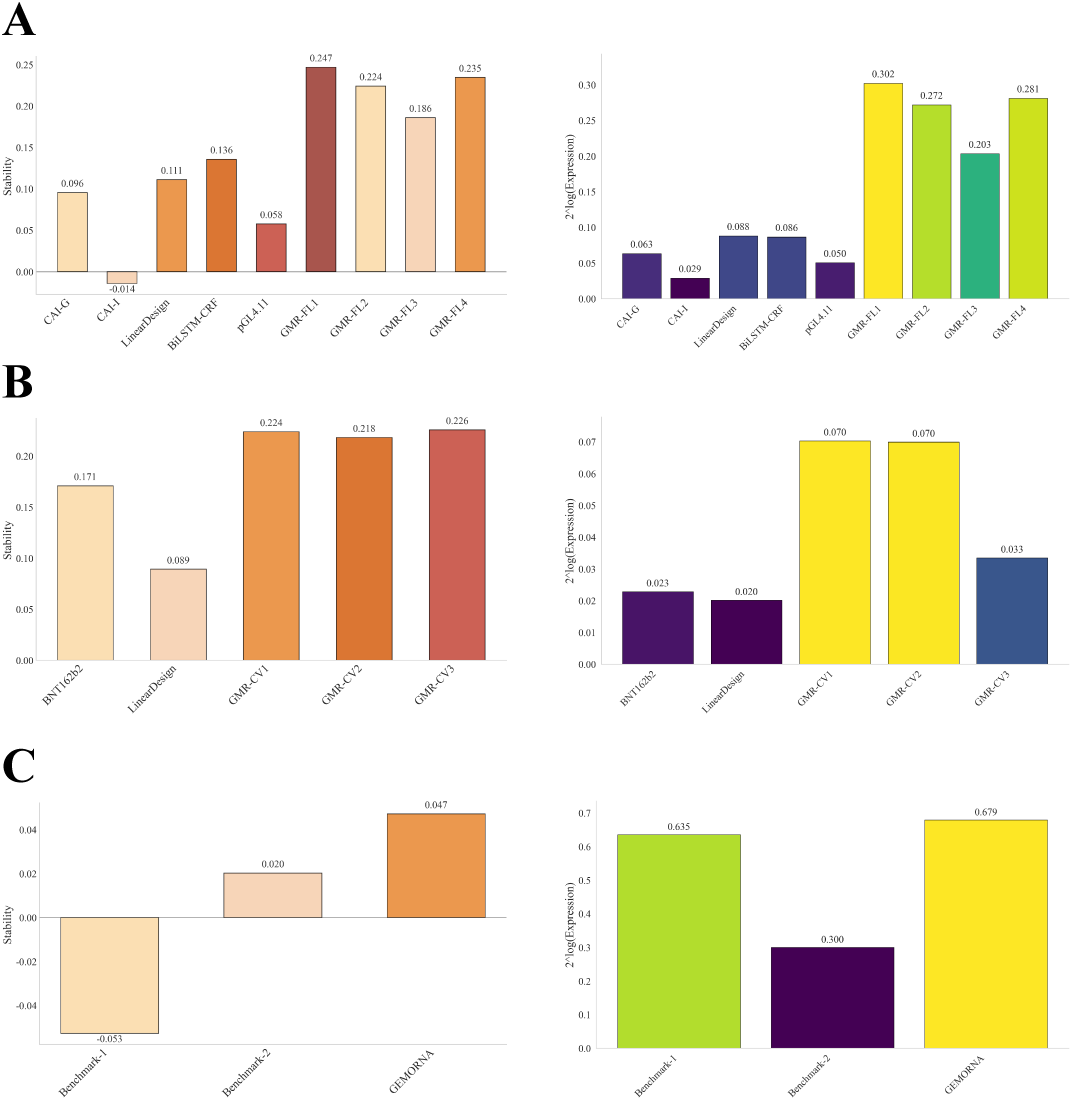
Evaluation of mRNA coding sequence encoding for firefly luciferase (A), Spike protein (B), and CD19 CAR (C) by using our stability and expression predictor.

## Notes

### Competing Interest Statement

The authors have declared no competing interest.

### Summary of Updates

Figure 3 revised; Figure 4 added; Figure 5 revised; additional author added; Supplemental files updated; Result 2.4 revised.

